# Transcriptional signatures of invasiveness in *Meloidogyne incognita* populations from sub-Saharan Africa

**DOI:** 10.1101/570770

**Authors:** Deborah Cox, Brian Reilly, Neil D. Warnock, Steven Dyer, Matthew Sturrock, Laura Cortada, Danny Coyne, Aaron G. Maule, Johnathan J. Dalzell

**Affiliations:** School of Biological Sciences, Queen’s University Belfast; International Institute of Tropical Agriculture, Kenya

**Author notes:** Joint first authors (Deborah Cox, Brian Reilly, Neil D. Warnock).

**Keywords:** Root-knot nematode, behaviour, invasion, transcriptome, microRNAs, plant parasitic nematode

## Abstract

*Meloidogyne incognita* is an economically important plant parasitic nematode. Here we demonstrate substantial variation in the invasiveness of four *M. incognita* populations relative to tomato. Infective (J2) stage transcriptomes reveal significant variation in the expression of protein-coding and non-coding RNAs between populations. We identify 33 gene expression markers (GEMs) that correlate with invasiveness, and which map to genes with predicted roles in host-finding and invasion, including neuropeptides, ion channels, GPCRs, cell wall-degrading enzymes and microRNAs. These data demonstrate a surprising diversity in microRNA complements between populations, and identify GEMs for invasiveness of *M. incognita* for the first time.

---

*Meloidogyne incognita* is a globally distributed and highly polyphagous parasite of crop plants (Coyne et al., 2018; Trudgill & Blok, 2001), demonstrating a surprisingly high level of adaptive variability for an asexual organism (Szitenberg et al., 2017). This adaptability is thought to play a role in the pests’ ability to rapidly evade sources of crop resistance. Consequently, *M. incognita* is becoming increasingly problematic, and current approaches to control are insufficiently robust or durable to provide reliable protection in the field (Davies & Elling, 2015). The natural variation between *M. incognita* populations is poorly understood (Bucki et al., 2017). This constitutes a substantial gap in our knowledge, which could hinder our ability to develop sources of durable resistance to field populations. The relatively high work burden of population maintenance in the laboratory, and inevitable domestication of *M. incognita* populations makes the assessment of field relevant inter-species variation a significant and ongoing technical challenge. In addition, access to populations that are native to Nagoya protocol (https://www.cbd.int/abs/) signatories can be problematic. Whilst the Nagoya protocol aims to promote equitable commercial outcomes arising from native genetic resources, opportunities for collaboration and extended sharing of resources are limited. Collectively, these challenges promote an artificial over-reliance on highly domesticated legacy strains, which are unlikely to reflect the genotypic or phenotypic spectra of field populations.

Although there are many potential approaches to developing crop parasite resistance, an improved understanding of parasite host-finding and invasion may facilitate the development of new strategies that prevent infection. This is preferable to sources of resistance that are active *in planta*, as it limits the opportunity for secondary pathogen infection, and minimises the metabolic burden of mounting a defence response to invading parasites. In this study, we assessed the host-finding and invasion behaviour of *M. incognita* populations that had been recently collected from field sites in Kenya and Nigeria, with the aim to relate observed behavioural variation to gene expression signatures using transcriptomic correlation. These data would improve our understanding of the link between genotype and phenotype, which may enable us to identify new targets for nematicide development, or biotechnological intervention.

We considered three populations collected from Nigeria, named Nig_1, Nig_2, and Nig_3, and one population from Kenya, named Ken_1. Our data demonstrate statistically significant variation in the propensity of these *M. incognita* populations to invade tomato cv. Moneymaker seedlings. Nig_1 is the most invasive, with a mean of 50.06 ±7.9 J2s (from a total of 200 J2s) invading within 24 h, followed by Ken_1 with a mean of 39.9 ±6.9, Nig_2 with a mean 27.3 ±7.6, and Nig_3 being the least invasive, with a mean of 2.5 ±0.8 J2s invading within 24 h (Figure 1).

**Figure 1.**
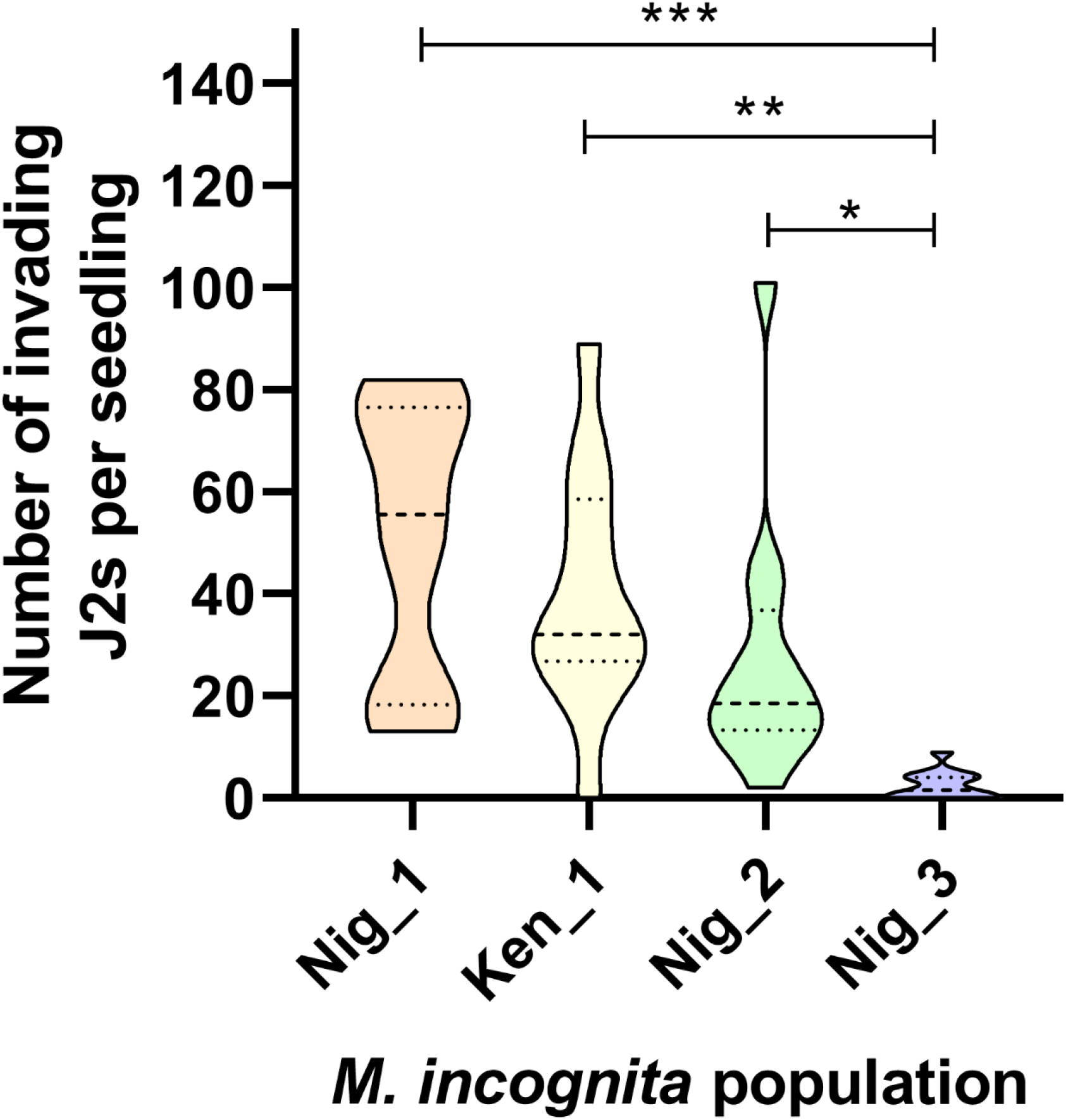
*Meloidogyne incognita* invasion of tomato cv. Moneymaker seedlings is highly variable. Violin plot showing number of J2s invading tomato seedlings, 24 h post exposure. Dashed lines indicate the median, dotted lines indicate the quartiles. Data assessed by ANOVA and Tukey’s multiple comparison test using Graphpad Prism 8; P<0.05*, P<0.01**, <0.001***. *M. incognita* populations were collected from field sites in Kenya and Nigeria. They were cultured on tomato cv. Moneymaker, in plant growth cabinets at 23°C, with a regular 16 h light, 8 h dark cycle for no more than two generations following field collection. Tomato seedling infection assays were conducted as in Warnock et al. (2016), using 200 J2s per seedling, inoculated into an agar slurry containing the tomato seedling.

We conducted high-throughput sequencing of protein-coding and non-coding RNAs from the infective J2 stage of each *M. incognita* population to understand the molecular basis of behavioural variation. Our data revealed that up to 6,232 (13.7%) transcripts were significantly up-regulated (P<0.0001****) relative to the most invasive population, Nig_1, with up to 4,908 (10.8%) down-regulated across pairwise comparisons (Figure 2A; supplemental file S1). Using mirDeep2, we identified 192 precursor microRNA genes across the four *M. incognita* populations, relating to 144 predicted mature microRNAs in *M. incognita* Nig_1; 146 in Nig_2; 105 in Nig_3; and 176 in the Ken_1 population. This constitutes a surprising diversity in microRNA complement between populations of the same species, with Ken_1 representing the major outlier, with 44 predicted mature microRNAs unique to that population (supplemental file S2). By way of comparison, a similar analysis using the entomopathogenic nematode *Steinernema carpocapsae* revealed variation from 269 to 273 predicted mature microRNAs across three populations (Warnock et al., 2018). Up to 52 (27%) of the predicted and conserved *M. incognita* microRNA genes were significantly up-regulated (P<0.0001****) relative to the most invasive Nig_1 population, with up to 23 (12%) down-regulated across pairwise comparisons (Figure 2B).

**Figure 2.**
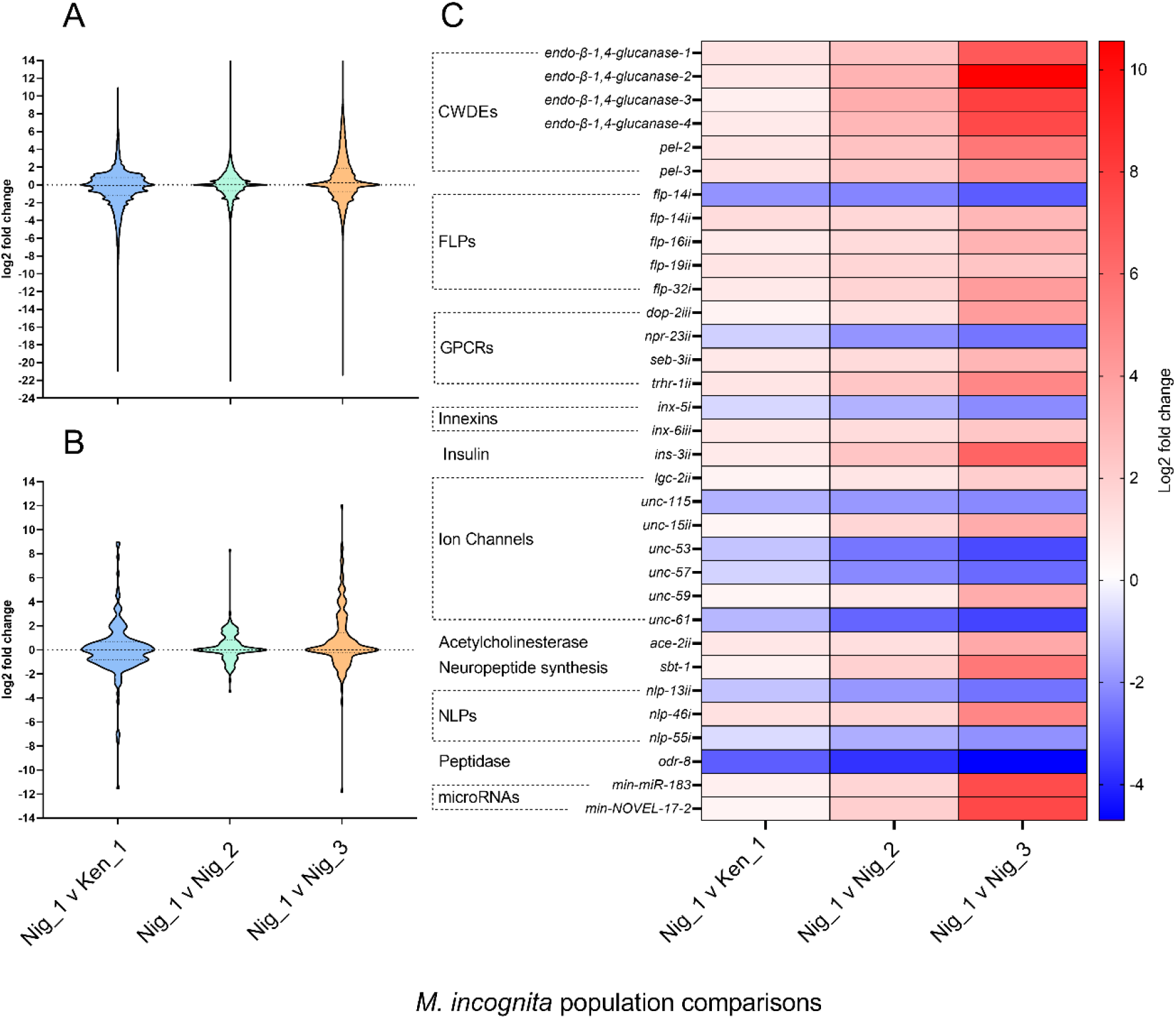
Transcriptomic signatures of *Meloidogyne incognita* invasiveness. Violin plots of log2 fold changes across pairwise population comparisons for (A) protein-coding genes and (B) microRNAs. (C) Summary heatmap of 33 selected GEMs, demonstrating the log2 fold change between pairwise comparisons, relative to the most invasive Nig_1 population. GEMs follow a gradient expression pattern that positively or negatively correlates with the population invasion phenotype; CWDEs – Cell Wall Degrading Enzymes. Figures were generated in Graphpad Prism 8. RNA extraction, library preparation, sequencing, bioinformatics and statistical analyses were conducted as in Warnock et al. (2018). Briefly, ~3000 J2s of each *M. incognita* population were used to extract total RNA, from which coupled 150 bp paired-end, and 50 bp single end illumina HiSeq libraries were prepared for each population, in triplicate. Libraries were sequenced on one illumina HiSeq 2500 lane. Following quality control, reads were mapped to the most recent *M. incognita* genome assembly (PRJEB8714, WBPS12, https://parasite.wormbase.org) using STAR and RSEM (Blanc-Mathieu et al., 2017; Howe et al., 2015; Dobin et al., 2013; Li et al., 2011). MicroRNAs were identified and quantified using MirDeep2 (Friedländer et al., 2012). Predicted microRNAs were named using a BLAST search against *C. elegans* microRNAs (www.miRBase.org), and previously identified *M. incognita* microRNAs (Zhang et al., 2015). Predicted microRNAs were named in line with *C. elegans* or *M. incognita* microRNAs represented as the top BLAST return, and if there was a sequence identity match greater than 80%. All novel *M. incognita* microRNAs were named sequentially, ensuring no overlap with names allocated to *C. elegans* or previously published *M. incognita* microRNAs. Predicted microRNA target genes were identified with MiRanda (Enright et al., 2003), using strict and unrestricted discovery modes, as in Warnock et al. (2018). Differentially expressed protein-coding and non-coding genes were identified using DESeq2 (Love et al., 2014). All sequencing datasets are available from the SRA database (Bioproject: PRJNA525879).

We populated a list of Gene Expression Markers (GEMs) that correlated, either positively or negatively, with the observed invasion phenotypes. This was achieved by arranging the population comparisons from most invasive to least invasive (Nig_1 vs Ken_1; Nig_1 vs Nig_2; Nig_1 vs Nig_3), and constraining gene lists to those that followed expression patterns consistent with the phenotypic trend. Correlating GEMs were identified when the log2 fold change quotients between adjacent comparisons were greater than one, with at least a P<0.05* difference between each population, and at least P<0.0001*** between the most and least invasive populations. Using this approach, we identified 485 GEMs that correlate with the observed invasion phenotype of *M. incognita*, comprising 483 protein-coding genes, and two microRNA genes; 242 GEMs correlate positively with the invasion phenotype, and 243 GEMs correlate negatively (Supplemental Files S1 & S2). On inspection of the invasion GEM list, we identified a total of 33 genes with predicted roles in the regulation of host-finding and invasion behaviour, including genes associated with the neuropeptidergic system, neuronal signalling, cell wall-degrading enzymes, and the two microRNA genes (Figure 2C). It is possible that other correlating genes play a functional role in the invasion phenotype, however we deemed that these 33 genes were most likely to exert the largest influence, based on known or predicted functionality.

Six neuropeptide genes correlated positively with *M. incognita* invasiveness, including *FMRFamide-like peptide 14ii (flp-14ii), flp-16ii, flp-19ii, flp-32i INSulin-like protein 3ii (ins-3ii)*, and *Neuropeptide-Like Protein 46i (nlp-46i).* Three neuropeptide genes, *flp-14i, nlp-13i* and *nlp-55i* were negatively correlated with the invasion phenotype (Figure 2C). Expression of a predicted prohormone convertase chaperone, *sbt-1*, which is required for the biosynthesis of neuropeptides in the free-living nematode *Caenorhabditis elegans* (Husson & Schoofs, 2007), also correlated positively with invasion behaviour. These data implicate the neuropeptidergic system, and FLPs in particular, in the modulation of *M. incognita* invasion behaviour. This corroborates previous observations of *flp* gene enrichment within the infective juvenile stage of many parasitic nematode species, and a role in the behavioural diversification of these stages (Lee et al., 2017). Indeed, our own work demonstrates similar associations between neuropeptidergic genes and the host-finding behaviour of *S. carpocapsae* (Warnock et al., 2018; Morris et al., 2017). Four putative neuropeptide G-Protein Coupled Receptor (GPCR) genes were also found to associate with *M. incognita* invasiveness, along with seven ion channel genes, two innexin genes, an acetylcholinesterase and a predicted *odr-8* peptidase homologue (Figure 2C). Within the 485 correlating GEMs, we also identified 62 novel genes, with no known function, or orthology to *C. elegans* genes (supplemental file S1). Six plant Cell Wall-Degrading Enzyme (CWDE) genes were also associated positively with invasion phenotypes, corresponding to four *endo-β-1,4-glucanase* genes, and two predicted pectate lyase *(pel)* genes (Figure 2C). Each CWDE gene is most highly expressed in the most invasive Nig_1 population, and display lowest expression in the least invasive population, consistent with a role in mediating the enzymatic degradation of the plant cell wall. If it can be demonstrated that certain CWDEs confer a specific advantage for the invasion of particular host species, it could point to new approaches to resistance based on the modification of cell wall composition, potentially in conjunction with recent developments in synthetic biology.

Non-coding microRNAs have been implicated in nematode behavioural variation (Warnock et al., 2018; Rauthan et al., 2017; reviewed in Ambros & Ruvkun, 2018), through the regulation of target gene expression. Our data reveal that the expression of two mature microRNAs correlates with the invasion phenotype of *M. incognita* populations (Figure 2C). Using miRanda to identify predicted gene targets, in both strict and unrestricted settings, reveals a surprising abundance, and interconnection between these microRNAs and neuropeptide genes, spanning the *flp, nlp* and *ins* families, in addition to GPCR and ion channel genes (supplemental file S3, S4). It has been suggested that microRNAs regulate developmental programmes through the coordinated and cooperative targeting of genes involved in specific biological functions (Zhang et al., 2009). Our data indicate that this may also be the case for behavioural regulation. For example, *Min-NOVEL-17-2* is predicted to simultaneously target: *flp-1i, flp-1ii, flp-11i, flp-11ii, flp-11iii, flp-33i, flp-33ii, flp-34i, ins-1i, ins-1ii, ins-1iv, ins-1v, ins-18i, nlp-8ii, nlp-12, nlp-13i, nlp-13ii, nlp-81iii*, in addition to a variety of other ion channel, GPCR and innexin genes (supplemental file S5). To further investigate the potential relationship between microRNAs and behavioural regulation, our analysis sought to identify predicted interactions that followed the expected trend for biologically interacting microRNA-mRNAs, at the level of mRNA abundance. A substantial literature has developed around microRNA induced mRNA decay in animals (reviewed in Iwakawa and Tomari, 2015), and our data demonstrate a correlation between numerous predicted microRNA targets, and expression patterns between populations, indicating that these microRNA-mRNA interactions may be biologically relevant. For example, *nlp-13ii* is identified as both an *in silico* predicted target of the novel microRNA *Min-NOVEL-17-2* and is demonstrated to follow an expression pattern consistent with microRNA-mediated mRNA decay across populations (Figure 3A). However, whilst *nlp-13i* is also a predicted target of *Min-Novel-17-2*, it does not follow an expression trend that is consistent with microRNA-mediated decay. This indicates either that the *nlp-13i* and *nlp-13ii* transcripts are expressed in different cells / tissues that only partially overlap with expression of *Min-NOVEL-17-2*, or that there are other transcript-specific features, which influence the tendency towards microRNA-mediated mRNA decay or translational inhibition. One possible explanation relates to altered secondary structure of UTR sequences, which may underpin differences in the bioavailability, or function of microRNA target sites. Based on *in silico* structural predictions using the Vienna RNAfold server (http://rna.tbi.univie.ac.at/), this does not appear to be a factor for the UTRs of *nlp-13* gene copies at least. Our analysis in Figure 3 focuses solely on predicted microRNA interactions with the 483 protein-coding genes identified as invasion GEMs, and on that basis makes no judgement on the likelihood of interactions with the many other predicted target genes listed above, which do not follow the stringency criteria used to populate the list of GEMs.

**Figure 3.**
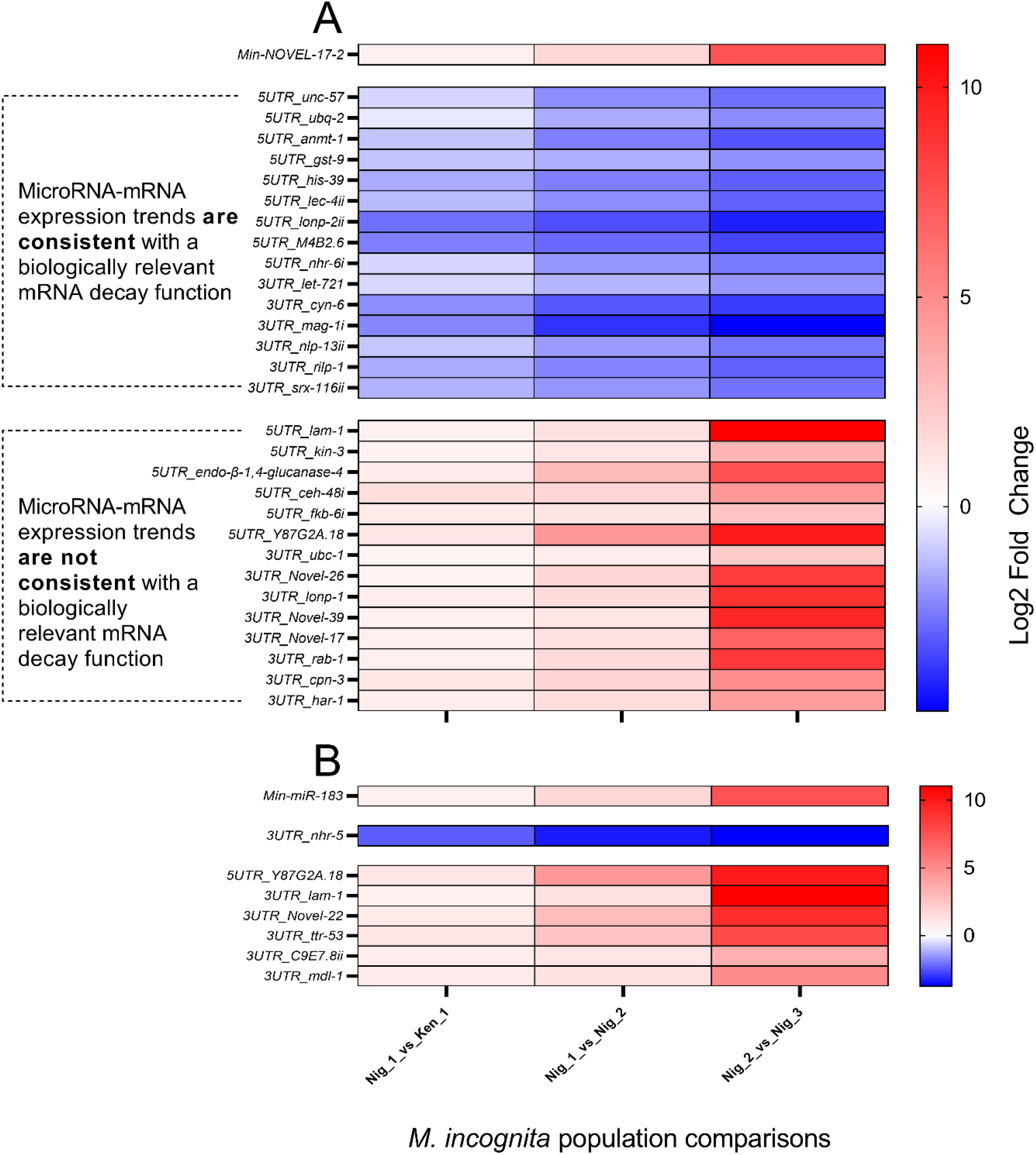
Differential expression of predicted microRNA targets suggests biologically relevant interactions across *Meloidogyne incognita* populations. (A) Heatmap demonstrating differential expression trends for the microRNA, *Min-NOVEL-17-2*, and predicted mRNA targets identified within the list of 483 protein-coding GEMs. The location of the predicted microRNA interaction point is indicated as a 5’UTR or 3’UTR suffix. 15 genes negatively correlate with *Min-NOVEL-17-2* changes across pairwise population comparisons, which suggests biologically relevant interactions, mediated through mRNA decay. 14 of the predicted targets, which follow the phenotype trend, do not correlate with *Min-Novel-17-2* in a way that suggests a biologically relevant interaction. (B) Heatmap demonstrating differential expression trends for the microRNA, *Min-miR-183* and predicted mRNA targets identified within the list of 483 protein-coding GEMs. Only one of the predicted target genes negatively correlates with microRNA differential expression trends, suggesting that the putative nuclear hormone receptor gene, *nhr-5*, is the only biologically relevant target within this set of invasion GEMs. The microRNA target analysis presented here is not intended to be an exhaustive treatment of all predicted mRNA interactions, focusing instead on the GEMs identified through transcriptomic and phenotypic correlation.

Apomictic *Meloidogyne* spp. are known to possess highly divergent hypotriploid genomes, with multiple variant gene copies (Szitenberg et al., 2017). The sequence variation of gene copies may reflect the functional diversification of common genetic elements for adaptive purposes, through hybridisation and selection. For example, a significant number of putative neuropeptide gene copies are found to encode the same predicted mature neuropeptide(s) within a variant mRNA sequence; such genes are identified within this manuscript by virtue of a Roman numeral suffix, assigned according to the order of discovery. Expression of *flp-14ii* correlates positively with increased invasiveness, whereas *flp-14i* correlates negatively with increased invasion behaviour (Figure 3, and Supplemental File S1). Whilst we have no insight to the relative role or function of gene copies, or if these copies co-localise, we have established that they can be differentially expressed, within and between populations of the same species. Our analysis of predicted microRNA interactions reveals a considerable amount of variation in the predicted 5’ and 3’ UnTranslated Regions (UTRs) of predicted neuropeptide gene copies, which underpins qualitative and quantitative variation in predicted microRNA targeting (supplemental files S3, S4 and S5). UTR sequence variation has received little attention in the literature for parasitic nematode species, however UTR sequences are known to be highly variable across developmental stages, and tissues of the model *C. elegans*, which drives the genic regulation of microRNA interactions (Blazie et al., 2010; Mangone et al., 2010). It is possible that the hypervariation of gene copy UTRs between *M. incognita* populations could be adaptive, driving functional divergence as a factor of differential microRNA targeting. Data support a similar hypothesis for UTR isoform variation and behavioural diversification of *S. carpocapsae* strains (Warnock et al., 2018). This could provide a functional explanation for the extraordinary variation and adaptiveness of apomictic *Meloidogyne* spp.

This study demonstrates a surprising behavioural variation amongst *M. incognita* populations that are native to Kenya and Nigeria and provides the first evidence of GEMs that correlate with the invasion phenotype. Furthermore, we observe substantial variation in the complement of microRNA genes between populations, and variation in gene UTR targets between variant gene copies, which could underpin behavioural adaptation to host and environment. These observations require detailed functional studies to ascertain the specific influence of implicated genes and microRNAs. Whilst the inevitable domestication of *M. incognita* populations under laboratory and greenhouse conditions constitutes a technical challenge for the study of field-relevant diversity and phenotype, we expect these populations to become better adapted to the experimental host. This could provide opportunity to track signatures of molecular adaptation over time, within an experimental evolutionary approach (reviewed by Kawecki et al., 2012).

## Acknowledgements

This project was funded by a Grand Challenges grant from the Bill and Melinda Gates Foundation, and a GCRF pilot grant from the Department for the Economy, Northern Ireland. We would like to thank Bastian Fromm (Stockholm University) for discussions around UTR secondary structure as a potential driver of microRNA interactions.

## Supporting information captions

Supplemental file S1. Global DESeq2 output across all pairwise population comparisons, and complete list of invasion GEMs.

Supplemental file S2. List of predicted microRNAs and global DESeq2 output across all pairwise population comparisons.

Supplemental file S3. MicroRNA target prediction analysis for global 5’UTRs.

Supplemental file S4. MicroRNA target prediction analysis for global 3’UTRs.

